# Marketable Yield and Grade Distribution of Sweetpotato (*Ipomoea batatas* [L.] Lam.) as Influenced by Production System and Season in a Temperate Climate

**DOI:** 10.64898/2026.10.14.759163

**Authors:** Luis O. Duque, Francesco Di Gioia

## Abstract

Protected culture systems offer a practical strategy for extending sweetpotato (*Ipomoea batatas* [L.] Lam.) production into temperate regions where cool soil temperatures and limited growing degree days (GDD) constrain storage root development. This study evaluated sweetpotato yield performance under four production systems (black plastic mulch [BPM], bare soil [BS], low tunnel [LT], and high tunnel [HT]) across two growing seasons (2022 and 2023) at Rock Springs, PA. Ten commercial cultivars were grown in a factorial randomized complete block design with four replications. Total yield (TY), marketable yield (MY), and individual grade yields (U.S. No. 1, Petite, Jumbo, Canner, and Cull) were assessed, along with seasonal soil temperatures (2023 only) and cumulative GDD (base 15.5°C). Year, Treatment, and their interaction were significant sources of variation for most yield components (P ≤ 0.05). In 2022, HT recorded the highest TY (21.1 Mg ha^−^¹) and BS the highest MY (16.8 Mg ha^−^¹), while LT produced the lowest values for both metrics. In 2023, BPM led all treatments in both TY (26.8 Mg ha^−^¹) and MY (26.6 Mg ha^−^¹). No single system consistently dominated across years, reflecting the importance of seasonal weather conditions on treatment response. Seasonal average soil temperatures in 2023 ranged from 20.7°C (BS) to 22.7°C (HT) but were not significantly correlated with grade yield at α = 0.05 (n = 4). Cumulative GDD ranged from 493 to 651 across treatments and years, well below the thresholds typical of major production regions. BPM provided the most consistent agronomic performance, supported by moisture conservation and weed suppression benefits that extended beyond the modest soil temperature advantage over bare soil. These results indicate that protected culture, particularly BPM, can support commercially viable sweetpotato production in temperate Pennsylvania, and that cultivar selection tailored to the production system is critical for optimizing yield and grade distribution.

## INTRODUCTION

Protected culture systems, ranging from simple low tunnels to fully managed high tunnels, are increasingly important in modern horticultural production. These structures extend the growing season and buffer crops against environmental stresses by moderating temperature extremes, reducing rainfall and pest pressure, and improving control over soil moisture and nutrient availability (Carey et al., 2009; Wells and Loy, 1993; Orzolek, 2013). Unlike fully controlled greenhouses, high- and low-tunnel systems are relatively low-cost and energy-efficient, making them suitable for small- and medium-sized farms and diversified specialty crop operations. Their growing adoption reflects a broader movement toward climate-resilient and sustainable production strategies that can maintain yield and quality under increasingly variable environmental conditions.

High tunnels are unheated, plastic-covered structures that create a semi-controlled environment by increasing air and soil temperatures, reducing evaporative losses, and excluding rainfall (Lamont, 2009). These conditions accelerate early growth, extend harvest windows, and improve crop uniformity and quality (Wells and Loy, 1993). Low tunnels, by contrast, are smaller, temporary coverings that provide short-term protection for transplants, often used in combination with mulches or drip irrigation to promote early establishment. Both systems modify the soil-plant microclimate, directly affecting plant physiology, phenology, and yield, particularly in temperate and transitional climates (Wells and Loy, 1993; Waterer, 1992). Although the benefits of protected culture are well documented for crops such as tomato (*Solanum lycopersicum*) (Maynard, 2025), cucumber (*Cucumis sativus*) (Rodriguez Izaba et al., 2021), and strawberry (*Fragaria × ananassa*) (Liu et al., 2024), their application to root and tuber crops, especially sweetpotato (*Ipomoea batatas* [L.] Lam.), remains limited.

In the United States, sweetpotato production is concentrated in southern states where warm-season conditions favor vegetative growth and storage root development, with North Carolina, California, Arkansas, Florida, Louisiana, and Mississippi accounting for more than 94% of planted area in 2018 (USDA-NASS, 2020). In contrast, production in the northeastern United States accounted for only about 1,300 acres (1.2% of national acreage) in 2017 (USDA-NASS, 2019), with most of the sweetpotatoes consumed in the region imported from southern states. Climatic constraints, including cool early-season soil temperatures, short frost-free periods, and limited growing degree days, restrict sweetpotato establishment and storage root bulking in northern environments. Nonetheless, increasing demand for locally grown, nutrient-dense crops has renewed interest in sweetpotato as a diversification option for small-scale and specialty crop systems in temperate zones.

Despite its tropical origin, sweetpotato is a resilient crop that performs across a wide range of soil types, climates, and management intensities (Jansson and Raman, 1991). Its adaptability has enabled cultivation from Florida to southern Ontario, Canada (Bouwkamp, 1985), and recent studies demonstrate that with appropriate cultivar selection and management, it can yield well in regions with shorter growing seasons and cooler temperatures (Sideman, 2015; Wees et al., 2015, 2016). In this context, protected culture systems offer a promising strategy to improve sweetpotato establishment and extend the effective growing season in northern production environments. However, quantitative data on the performance of different protected culture configurations, particularly their effects on soil microclimate, crop development, and marketable yield, are limited for northeastern U.S. conditions.

To address this gap, the present study evaluated sweetpotato performance under four production systems: high tunnel, low tunnel, black plastic mulch, and bare soil. Specifically, we (1) characterized soil temperature and accumulated heat units across production systems over two growing seasons; (2) assessed marketable yield and grade distribution; and (3) evaluated cultivar responses to protected culture environments.

## MATERIALS AND METHODS

### Site description

The study was conducted over two growing seasons (2022 and 2023) at the Russell E. Larson Agricultural Research Center, The Pennsylvania State University, Rock Springs, PA (40°42′38.1″ N, 77°57′52.1″ W). The site has a Murrill channery silt loam soil (fine-loamy, mixed, semiactive, mesic Typic Hapludult) with a 0-3% slope. Monthly temperature and precipitation data were obtained from the Network for Environmental and Weather Applications (NEWA) station in Rock Springs, PA (40°42′32.5″ N, 77°57′06.9″ W). The preceding crop in both years was winter wheat (*Triticum aestivum* L.) grown as a cover crop.

Field soil samples (black plastic mulch, bare soil, and low tunnel plots) were collected 15 days before field preparation each year and analyzed by the Agricultural Analytical Services Laboratory, The Pennsylvania State University, University Park, PA. Soil test results indicated phosphorus (as P□O□) and potassium (as K□O) concentrations of 36.6 and 240 kg ha^−^¹, respectively, as determined by the Mehlich 3 extraction method. Prior to treatment establishment, 252 kg ha^−^¹ P and 146 kg ha^−^¹ K were incorporated to meet sweetpotato fertilization recommendations, and 84 kg ha^−^¹ calcium nitrate (15.5-0-0) was applied preplant based on soil test results. An additional 84 kg ha^−^¹ calcium nitrate was applied through fertigation to all treatments 30 days after transplanting, corresponding to the onset of storage root initiation.

The high tunnel was located on a contiguous but separate field, and soil tests from that area indicated P□O□ and K□O concentrations of 282 and 446 kg ha^−^¹, respectively. Prior to treatment establishment, 118 kg ha^−^¹ P and no supplemental K were incorporated to meet sweetpotato fertilization recommendations. Preplant and fertigation calcium nitrate applications followed the same protocol as the other treatments. The fertilization regime was identical in both years across all treatments.

All plots were drip irrigated as needed using the feel-and-appearance method (USDA-NRCS, 1998). Weeds between rows were manually removed from June through August; thereafter, vine growth provided full canopy closure, suppressing further weed emergence.

### Sweetpotato cultivars

All sweetpotato slips used in both years were G2 planting stock provided by K. V. Pecota, North Carolina State University, Raleigh, NC (Table 1).

**Table 1.** Origin, skin color, and flesh color of commercial sweetpotato cultivars evaluated at Rock Springs, PA, in 2022 and 2023.

| Cultivar | Source | Skin Color | Flesh Color | Yr(s) used | Citation |
| --- | --- | --- | --- | --- | --- |
| Averre | NCSU | Orange/Copper | Orange | 2022, 2023 | Yencho et al., 2018 |
| Beauregard | LSU | Orange/Copper | Orange | 2022, 2023 | Rolston et al., 1987 |
| Bayou Belle | LSU | Red | Orange | 2022, 2023 | LaBonte et al., 2013 |
| Carolina Ruby | NCSU | Red | Orange | 2023 | Collins et al., 1999 |
| Covington | NCSU | Orange/Copper | Orange | 2022, 2023 | Yencho et al., 2008 |
| Hatteras | NCSU | Rose/Copper | Orange | 2022 | Schultheis, et al., 2013 |
| Murasaki | LSU | Purple | White | 2023 | LaBonte et al., 2008 |
| O' Henry | Henry Wayne Bailey | Copper | White | 2022, 2023 | LaBonte et al., 2011 |
| Purple Splendor | NCSU | Dark Purple | Purple | 2023 | Yencho et al., 2022 |
| White Bonita | LSU | Tan | White | 2022, 2023 | LaBonte et al., 2011 |

### Treatments

#### Black plastic mulch

Raised beds were formed using a combined raised-bed, mulch, and drip-tape layer (Model 2400 Mini Layer; Rain-Flo Irrigation, East Earl, PA). Beds were 0.91 m wide and 0.20 m high, with 1.52 m between bed centers. Each bed was covered with 25-µm black embossed plastic mulch (part no. BLK324; Rain-Flo Irrigation), and a single line of drip irrigation tape (T-Tape; T Systems International, San Diego, CA) with 20-cm emitter spacing was installed beneath the mulch. Sweetpotato slips were hand-planted in a single row at 30-cm in-row spacing on 15 June 2022 and 15 June 2023. The experiment was arranged in a randomized complete block design with four replicates per treatment; each plot consisted of 10 plants separated by 1.52-m alleys.

#### Bare soil

A bare-soil treatment (no plastic mulch) was established on flat, tractor-disked ground. Rows were spaced 0.91 m center-to-center, and a single drip tape line with 20-cm emitter spacing was laid along the soil surface at the center of each row. Slips were hand-planted at 30-cm in-row spacing on the same dates as the black plastic mulch treatment. This system is representative of conventional regional sweetpotato production practices.

#### Low tunnel

Raised beds were constructed as described for the black plastic mulch treatment (0.91 m wide, 0.20 m high, 1.52 m center-to-center) but without plastic mulch. A single drip tape line with 20-cm emitter spacing was installed along the center of each bed. Retractable low tunnels (Bio Plus system; Dubois Agrinovation, Saint-Rémi, QC, Canada; PK TFLEX50) were covered with clear polyethylene film (38 µm). Sweetpotato slips were hand-planted at 30-cm in-row spacing on 27 May 2022 and 3 May 2023. Low tunnels were removed 21 days after planting to permit adequate ventilation and prevent excessive internal heat, humidity, and vine contact with the film. The earlier planting date afforded by the low tunnel resulted in an extended growing season relative to the black plastic mulch and bare soil treatments.

#### High tunnel

A fourth treatment was established on bare soil under a stationary high tunnel measuring 15 m long × 9 m wide × 3 m tall, covered with polyethylene greenhouse film (Tufflite IV; product ID 7301; Johnny’s Selected Seeds, Fairfield, ME). Rows were spaced 0.91 m center-to-center with a single drip tape line per row. End walls were rolled up for ventilation in response to ambient temperature; the structure was not moved during the experiment. Sweetpotato slips were hand-planted at 30-cm in-row spacing on 13 May 2022 and 17 May 2023.

### Harvest and grading

Sweetpotatoes were harvested on 20 Sept 2022 and 22 Sept 2023 (high tunnel; 121 and 128 days after transplanting), 4 Oct 2022 and 28 Sept 2023 (low tunnel; 131 and 121 days after transplanting), and 20 Oct 2022 and 13 Oct 2023 (black plastic mulch and bare soil; 127 and 121 days after transplanting). Vines were cut with a walk-behind string trimmer (model ST 100; Cub Cadet, Cleveland, OH) and plastic mulch was removed by hand. Storage roots were lifted using a one-row potato digger (model D-10M; U.S. Small Farm Equipment Co., Worland, WY) and graded according to U.S. standards for sweetpotatoes (USDA Agricultural Marketing Service, 2026) into the following categories: U.S. No. 1 (diameter 4.4-8.9 cm; length 7.6-22.9 cm), Petite (diameter 3.8-5.7 cm; length 7.6-17.8 cm), Jumbo (diameter >8.9 cm or length >22.9 cm), Canner (diameter ≤5.7 cm or length ≤7.6 cm, no external damage), and Cull (any size with off-shapes or damage from bruising, scarring, growth cracks, decay, insects, or other causes).

Marketable yield (MY) was calculated as the sum of all grades excluding culls; total yield (TY) included all grades. All storage roots were counted, measured, and weighed. Data were tested for normality and analyzed using analysis of variance (ANOVA) in JMP Student (version 19; SAS Institute, Cary, NC). When the overall F-test was significant (P ≤ 0.05), means were separated using Tukey’s honest significant difference test (P ≤ 0.05).

### Growing degree days

Daily growing degree days (GDD) were calculated as:

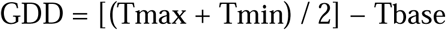

where Tmax and Tmin are the daily maximum and minimum air temperatures (°C), and Tbase is the base temperature set to 15.5°C, which represents the minimum temperature threshold for active sweetpotato growth (Jansson and Raman, 1991; Bouwkamp, 1985). When the daily mean air temperature fell below the base temperature, the GDD was set to zero. Cumulative GDD was calculated as the sum of daily values from the transplanting date to harvest for each treatment and growing season.

### Soil temperature monitoring

Soil temperature was continuously monitored throughout the 2023 growing season across all four production systems. Data were not collected during the 2022 growing season; therefore, all soil temperature analyses presented here are based exclusively on 2023 observations.

In the black plastic mulch, bare soil, and low tunnel treatments, soil temperature was recorded using HOBO Bluetooth pendant temperature/light data loggers (model MX2202; Onset Computer Corporation, Bourne, MA). Sensors were inserted at a depth of 12 cm within the plant row at the time of transplanting and programmed to record at 15-min intervals. In the low tunnel treatment, the sensor captured the thermal microenvironment created by the tunnel during the initial 21 days after transplanting; after tunnel removal, in-row temperature was monitored for the remainder of the season. Data were retrieved wirelessly via Bluetooth using the HOBOconnect mobile application (v1.6.1.1400; Onset Computer Corporation).

In the high tunnel, soil temperature was recorded using a 5TM soil moisture and temperature sensor (Meter Group, Pullman, WA) connected to an EM50 data logger (Decagon Devices, Pullman, WA). The sensor was inserted at 12 cm depth within the plant row, consistent with the other treatments, and data were recorded at 10-min intervals.

Raw interval data from all treatments were aggregated to daily mean, maximum, and minimum values for analysis and graphical presentation. Observations recorded after harvest, when sensor disturbance produced erratic values, were excluded from the analysis.

## RESULTS

### Weather conditions at Rock Springs, PA (2022 and 2023)

Air temperature, precipitation, and growing degree day (GDD; base 15.5°C) accumulation during the 2022 and 2023 growing seasons (May-October) at Rock Springs, PA, are presented in Fig. 1A, 1B, and 1C. These data contextualize environmental conditions during the experimental period and highlight interannual climatic variability.

**Figure 1.**
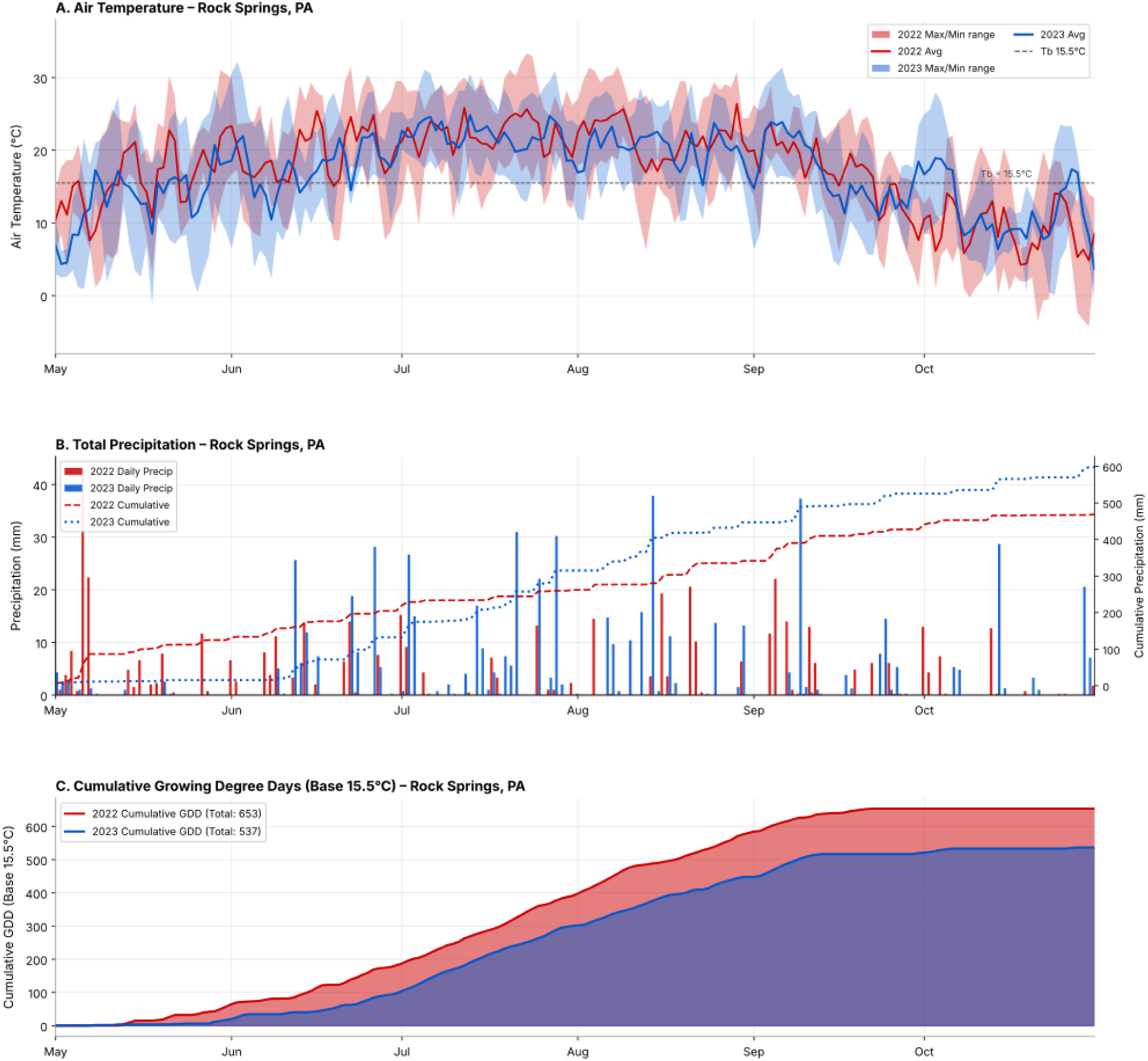
Daily and cumulative weather conditions recorded at Rock Springs, PA, during the 2022 and 2023 sweetpotato growing seasons (May–October). (A) Air Temperature (°C): Solid lines represent daily average air temperature for 2022 (orange) and 2023 (blue). Dashed lines indicate daily maximum temperatures; dotted lines indicate daily minimum temperatures. Shaded areas represent the daily min–max temperature range for each year. The horizontal dashed grey line indicates the base temperature (15.5°C) used in growing degree-day calculations. (B) Total and Cumulative Precipitation (mm): Bars represent daily total precipitation for 2022 (red) and 2023 (blue), displayed side by side for each date. Dotted lines on the secondary y-axis (right) represent cumulative seasonal precipitation for each year. (C) Cumulative Growing Degree Days (GDD, Base 15.5°C): Lines represent cumulative GDD accumulation throughout the season for 2022 (orange, total = 653 GDD) and 2023 (blue, total = 533 GDD). Shaded areas below each line indicate the magnitude of GDD accumulation over time.

In 2022, the seasonal mean air temperature was 18.1°C, with a peak daily maximum of 33.3°C on 23 July and a minimum of −0.3°C on 9 Oct. Monthly average temperatures followed a unimodal pattern, peaking in July (22.2°C) and declining through September (16.9°C) and October (9.3°C). Total seasonal precipitation was 466.6 mm over 73 rain days, with May receiving the highest monthly accumulation (125.0 mm), including a single-day maximum of 43.2 mm on 6 May. Precipitation from June through September ranged from 57.4 mm (July) to 86.4 mm (September). Cumulative GDD totaled 653.4 for the season, with 523.0 GDD accumulated between June and August.

The 2023 season had a slightly cooler seasonal mean temperature (17.3°C), with temperatures peaking similarly in July (22.1°C). The season began cooler, however, with a May average of 13.3°C and a seasonal minimum of −1.3°C on 18 May. Total precipitation was higher in 2023 (569.0 mm), driven by above-average rainfall in June (117.1 mm), July (182.6 mm), and August (131.8 mm); the single-day maximum was 37.8 mm on 14 Aug. May 2023 was exceptionally dry, receiving only 15.5 mm. Cumulative GDD in 2023 totaled 533.2, approximately 120 GDD fewer than in 2022, reflecting cooler conditions in May and August. Together, 2022 was warmer and more thermally productive while 2023 was wetter and thermally limited, providing important context for interpreting treatment differences in sweetpotato yield across the two growing seasons.

### Yield performance across treatments and years

Total yield (TY), marketable yield (MY), and individual grade yields (U.S. No. 1, Petite, Jumbo, Canner, and Cull) were assessed across four treatments: black plastic mulch (BPM), bare soil (BS), high tunnel (HT), and low tunnel (LT), over two growing seasons (2022 and 2023). The ANOVA indicated significant main effects of Year (Y) and Treatment (T) on TY and MY, as well as significant Y × T interactions for most grade components (Table 2). Total and marketable yield results are presented in Fig. 2A. In 2022, HT recorded the highest TY (21.1 Mg ha^−^¹) and BS the highest MY (16.8 Mg ha^−^¹), while LT recorded the lowest values for both metrics (TY: 6.4 Mg ha^−^¹; MY: 5.1 Mg ha^−^¹). In 2023, BPM was the top-yielding treatment for both TY (26.8 Mg ha^−^¹) and MY (26.6 Mg ha^−^¹), representing the largest between-season gain of any treatment. LT improved substantially from 2022 to 2023 (TY: 6.4 to 21.2 Mg ha^−^¹; MY: 5.1 to 19.4 Mg ha^−^¹), while BS declined in TY (19.7 to 16.4 Mg ha^−^¹) but maintained similar MY (16.8 to 16.2 Mg ha^−^¹).

**Figure 2.**
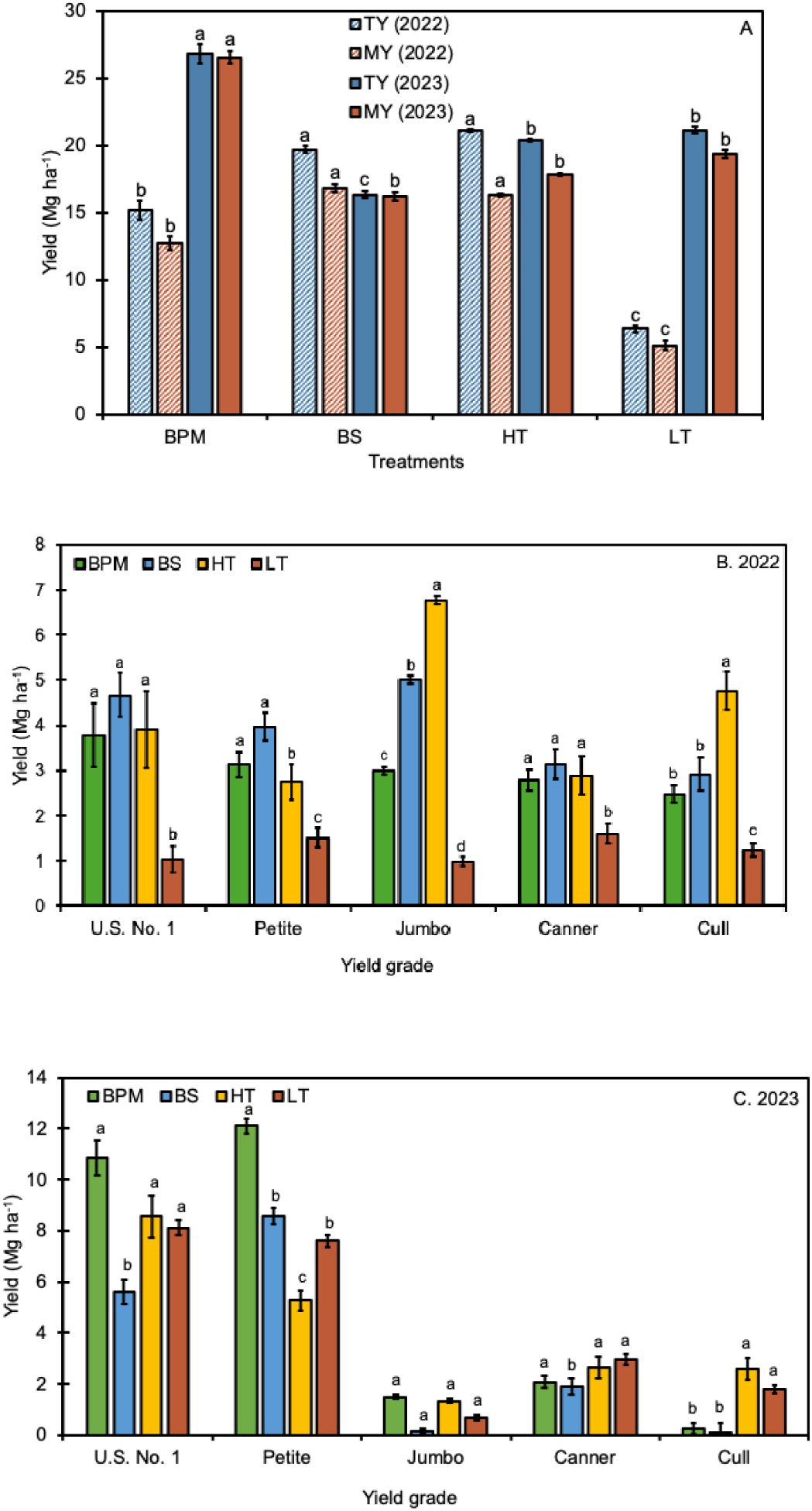
Sweetpotato yield by production system across two growing seasons and yield grades in southeastern Pennsylvania. (A) Total yield (TY) and marketable yield (MY) by production system in 2022 and 2023. Hatched bars represent 2022; solid bars represent 2023. (B) Yield by grade and production system in 2022. (C) Yield by grade and production system in 2023. Values are means ± 1 SE (n = 4 except high tunnel in 2022, n = 3). Within each yield category or grade, bars sharing the same lowercase letter do not differ significantly (Tukey HSD, *P* ≤ 0.05); letters are not comparable across yield categories or grades. No significant treatment differences were detected for Jumbo yield in 2023 (*P* > 0.05). Grade definitions follow USDA standards: U.S. No. 1 = 4.4–8.9 cm diameter; Petite < 4.4 cm diameter; Jumbo > 8.9 cm diameter; Canner = irregular or slightly defective roots meeting minimum size; Cull = unmarketable. BPM = black plastic mulch; BS = bare soil; HT = high tunnel; LT = low tunnel.

**Table 2.** Type III analysis of variance for sweetpotato yield by grade in a factorial randomized complete block design across two growing seasons (2022 and 2023) at Rock Springs, PA.

| Source of variation | df | TY |  | MY |  | U.S. No. 1 |  | Petite |  | Jumbo |  | Canner |  | Cull |  |
| --- | --- | --- | --- | --- | --- | --- | --- | --- | --- | --- | --- | --- | --- | --- | --- |
|  |  | F | Sig. | F | Sig. | F | Sig. | F | Sig. | F | Sig. | F | Sig. | F | Sig. |
| Year (Y) | 1 | 0.02 | NS | 0.01 | NS | 0.13 | NS | 0.04 | NS | 0.00 | NS | 0.67 | NS | 0.25 | NS |
| Treatment (T) | 3 | 11.84 | *** | 8.29 | *** | 1.93 | NS | 2.30 | NS | 17.07 | *** | 4.43 | ** | 17.65 | *** |
| Cultivar (C) | 9 | 0.22 | NS | 0.28 | NS | 0.31 | NS | 0.19 | NS | 0.00 | NS | 0.01 | NS | 0.01 | NS |
| Y × T | 3 | 11.47 | *** | 8.01 | *** | 4.41 | ** | 8.28 | *** | 11.24 | *** | 8.02 | *** | 15.21 | *** |
| Y × C | 9 | 0.31 | NS | 0.27 | NS | 0.17 | NS | 0.12 | NS | 0.00 | NS | 0.73 | NS | 0.22 | NS |
| T × C | 27 | 0.31 | NS | 0.27 | NS | 0.17 | NS | 0.12 | NS | 0.00 | NS | 0.73 | NS | 0.22 | NS |
| Y × T × C | 27 | 0.31 | NS | 0.27 | NS | 0.17 | NS | 0.12 | NS | 0.00 | NS | 0.73 | NS | 0.22 | NS |
| Residual | 165 | — | — | — | — | — | — | — | — | — | — | — | — | — | — |

Yield components include total yield (TY), marketable yield (MY), and five USDA grade classes: U.S. No. 1 (4.4–8.9 cm diameter), Petite (< 4.4 cm diameter), Jumbo (> 8.9 cm diameter), Canner (irregular or slightly defective roots meeting minimum size), and Cull (unmarketable). F-statistics were estimated using ordinary least squares with block nested within year [Block (Year)] included as a covariate to account for the unbalanced design (high tunnel, 2022: $n = 3$ ; all other treatment-year combinations: $n = 4$ ). Degrees of freedom: Year (Y) = 1; Treatment (T) = 3; Cultivar (C) = 3; Y × C = 3; T × C = 9; Y × T × C = 27; Residual = 165. NS = not significant; \* $P \leq 0.05$ ; \*\* $P \leq 0.01$ ; \*\*\* $P \leq 0.001$ .

Grade yield distributions are presented in Fig. 2B and 2C. U.S. No. 1 yield was highest under BS in 2022 (4.7 Mg ha^−^¹) but increased substantially across all treatments in 2023, with BPM recording the highest value (10.9 Mg ha^−^¹). Petite yield followed a similar pattern, with BS leading in 2022 (4.0 Mg ha^−^¹) and BPM in 2023 (12.1 Mg ha^−^¹). Jumbo yield was notably higher in 2022 than in 2023 across all treatments, with HT producing the most in 2022 (6.8 Mg ha^−^¹) and BPM the most in 2023 (1.5 Mg ha^−^¹); no significant treatment differences were detected for Jumbo yield in 2023 (Table 2). Canner yield was relatively consistent across treatments and seasons, ranging from 1.6 to 3.2 Mg ha^−^¹ in 2022 and 1.9 to 3.0 Mg ha^−^¹ in 2023. Cull yield was highest under HT in both seasons (4.8 and 2.6 Mg ha^−^¹ in 2022 and 2023, respectively); all other treatments showed reduced Cull fractions in 2023 relative to 2022. Overall, 2023 was characterized by substantially higher U.S. No. 1 and Petite yields and lower Jumbo and Cull fractions across all treatments compared with 2022, indicating a shift toward premium-grade production in the second season.

### Cultivar performance across treatments and years

Cultivar marginal means pooled across years and treatments are presented in Table 3. Yield performance varied considerably across cultivars, treatments, and growing seasons. In 2022, the highest TY values were recorded under HT, with ‘Purple Splendor’ (28.5 Mg ha^−^¹) and ‘Bayou Belle’ (28.2 Mg ha^−^¹) as the top-performing cultivars. Under BS, ‘Hatteras’ achieved the highest TY (25.5 Mg ha^−^¹) and MY (22.6 Mg ha^−^¹) of any treatment-cultivar combination in 2022. The BPM treatment yielded the highest TY for ‘Averre’ (20.0 Mg ha^−^¹), while LT consistently produced the lowest TY and MY across cultivars, with ‘White Bonita’ recording the season minimum (TY: 2.5 Mg ha^−^¹). HT was characterized by high Jumbo fractions, peaking at 7.4 Mg ha^−^¹ for ‘Murasaki’ and 7.3 Mg ha^−^¹ for ‘Bayou Belle’, and elevated Cull yields, particularly for ‘Bayou Belle’ (8.6 Mg ha^−^¹) and ‘Beauregard’ (5.2 Mg ha^−^¹). U.S. No. 1 yields in 2022 were generally modest across all treatments, with BS-‘Beauregard’ (7.2 Mg ha^−^¹) and BS-‘Hatteras’ (7.0 Mg ha^−^¹) recording the highest values.

**Table 3.**
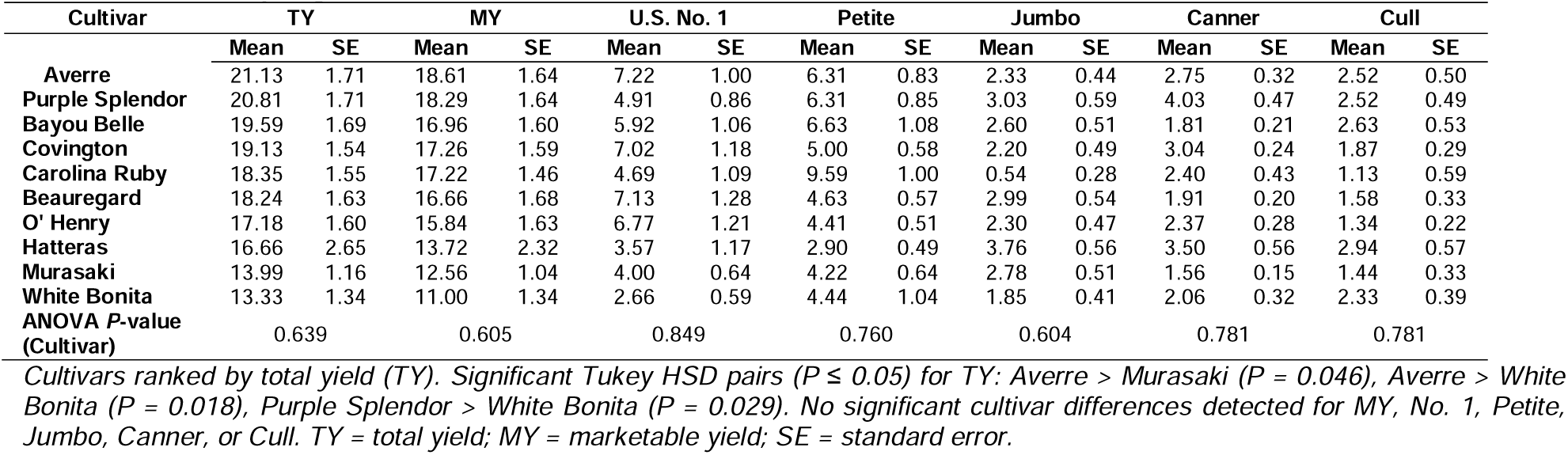
Sweetpotato cultivar marginal means for yield grade components (Mg ha^⁻¹^ ± SE), pooled across years and treatments. Rock Springs, PA, 2022–2023.

In 2023, BPM was the highest-yielding treatment across most cultivars. ‘O’Henry’ under BPM achieved the highest TY (30.9 Mg ha^−^¹) and MY (30.8 Mg ha^−^¹) of any treatment-cultivar combination across both years, with a U.S. No. 1 yield of 18.4 Mg ha^−^¹. ‘Beauregard’ and ‘Averre’ under BPM also recorded high TY values of 30.6 and 30.0 Mg ha^−^¹, respectively. Under LT, ‘Covington’ produced the highest TY (31.4 Mg ha^−^¹) and MY (29.8 Mg ha^−^¹) within that treatment, with a U.S. No. 1 yield of 17.4 Mg ha^−^¹. ‘Purple Splendor’ performed notably well under HT in 2023 (TY: 29.9 Mg ha^−^¹; MY: 28.4 Mg ha^−^¹). ‘Murasaki’ underperformed in 2023, recording the lowest TY under both BPM (19.0 Mg ha^−^¹) and LT (6.0 Mg ha^−^¹). Cull fractions were markedly reduced under BPM and BS in 2023, with several combinations recording zero Cull yield. Cultivar marginal means for all yield grade components are summarized in Table 3.

### Cumulative growing degree day and yield relationships

Cumulative GDD (base 15.5°C) from transplanting to harvest varied between years and among treatments (Fig. 1 and 3). In 2022, GDD accumulation ranged from 555.4 (BPM and BS) to 650.5 (HT), reflecting earlier transplanting dates for HT (13 May) and LT (27 May) compared with BPM and BS (15 June). In 2023, GDD values were lower overall and less variable across treatments, ranging from 493.0 (BPM and BS) to 516.6 (LT), consistent with a cooler early season.

**Figure 3.**
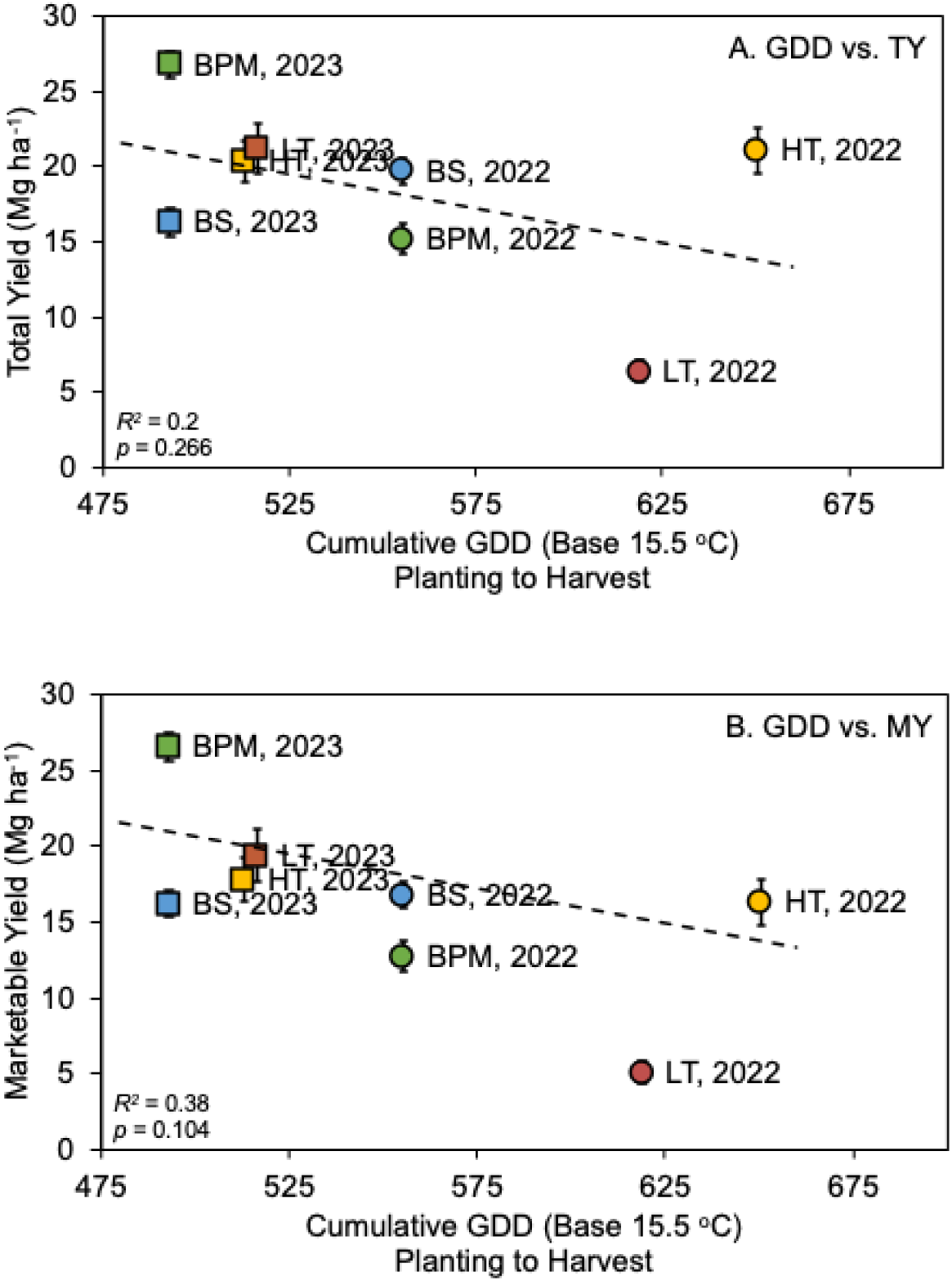
Relationship between cumulative growing degree days (GDD, base 15.5°C) accumulated from transplanting to harvest and sweetpotato yield across four protected culture treatments at Rock Springs, PA, in 2022 and 2023. (A) Total Yield (TY): Each point represents the mean total yield (Mg ha^⁻^¹) for a given treatment and year — black plastic mulch (BPM, green), bare soil (BS, blue), high tunnel (HT, yellow), and low tunnel (LT, red). Circles (●) represent 2022 data; squares (▪) represent 2023 data. Vertical bars indicate standard errors of the mean. The dashed grey line represents the linear regression fit for the combined 2022–2023 dataset (r = −0.448, R² = 0.200, p = 0.266, n = 8). (B) Marketable Yield (MY): Each point represents the mean marketable yield (Mg ha^⁻^¹), calculated as the sum of U.S. No. 1, Petite, Jumbo, and Canner grades, excluding Cull. Circles (●) represent 2022 data; squares (▪) represent 2023 data. Vertical bars indicate standard errors of the mean. The dashed grey line represents the linear regression fit for the combined 2022–2023 dataset (r =−0.616, R² = 0.380, p = 0.104, n = 8).

When GDD and yield data from both seasons were combined (n = 8 treatment-year observations), the linear relationship between cumulative GDD and TY was negative and non-significant (Fig. 3A; combined R² = 0.20, P = 0.266). The relationship between GDD and MY was also negative and non-significant (Fig. 3B; combined R² = 0.38, P = 0.104). The absence of a positive GDD-yield relationship was largely attributable to the LT treatment in 2022, which accumulated the second-highest GDD (618.8) yet produced the lowest TY (6.4 Mg ha^−^¹) and MY (5.1 Mg ha^−^¹) of all treatments. These results indicate that cumulative GDD alone is an insufficient predictor of sweetpotato yield across protected culture systems at this location.

### Soil temperature and yield relationships

Soil temperature was monitored only during the 2023 growing season; data were not collected in 2022 (Fig. 4). Seasonal average soil temperatures ranged from 20.7°C (BS) to 22.7°C (HT), with BPM (21.3°C) and LT (21.3°C) recording intermediate values (Fig. 4). HT maintained the warmest and most stable soil temperatures throughout the season. BPM exhibited relatively stable daily temperatures with a narrower daily range than BS. In the LT treatment, soil temperatures were elevated relative to ambient conditions during the first 21 days after transplanting; following tunnel removal on 21 June 2023, temperatures converged toward values more consistent with BPM and BS.

**Figure 4.**
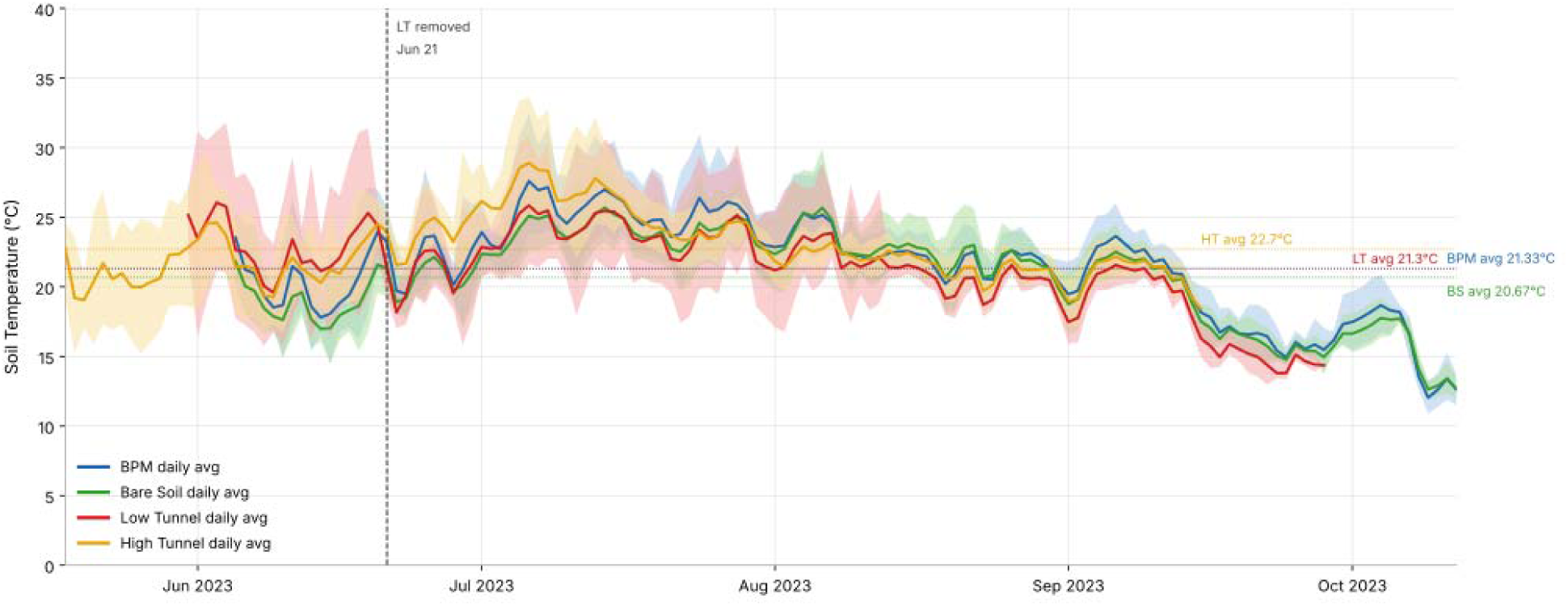
Daily soil temperature (°C) at 12 cm depth across four protected culture treatments during the 2023 growing season at Rock Springs, PA. Solid lines represent daily average soil temperature for black plastic mulch (BPM; green), bare soil (BS; blue), low tunnel (LT; red), and high tunnel (HT; yellow). Shaded bands indicate the daily minimum-to-maximum temperature range for each treatment. Dotted horizontal lines indicate the seasonal average soil temperature for each treatment (BPM: 21.3°C; BS: 20.7°C; LT: 21.3°C; HT: 22.7°C). The vertical dashed line marks the date of low tunnel removal (June 21, 2023). Soil temperature was recorded at 15-minute intervals using HOBO MX2202 pendant loggers (BPM, BS, and LT) and at 10-minute intervals using a Decagon EM50 datalogger equipped with 5TM soil moisture and temperature sensors (HT). Data are presented from transplanting through harvest for each treatment: BPM and BS through October 12, LT through September 28, and HT through September 15, 2023.

Despite recording the highest seasonal average soil temperature, HT did not produce the highest marketable yield in 2023. BPM produced the highest TY (26.8 Mg ha^−^¹) and MY (26.6 Mg ha^−^¹) despite recording only an intermediate seasonal soil temperature (21.3°C), while HT ranked third in both metrics (TY: 20.4 Mg ha^−^¹; MY: 17.8 Mg ha^−^¹). These results suggest that factors beyond soil thermal accumulation, including moisture retention under plastic mulch and reduced weed pressure, contributed substantially to yield outcomes.

Pearson correlation coefficients between seasonal average soil temperature and individual grade yields are summarized in Fig. 5F. Correlations were weak to moderate and non-significant at α = 0.05 (n = 4; Fig. 5A–F). U.S. No. 1 yield showed a weak positive trend (r = 0.393, R² = 0.155, P = 0.607), while Petite yield showed a moderate negative relationship (r = −0.601, R² = 0.361, P = 0.399), suggesting that cooler soils were associated with proportionally higher Petite root production. Jumbo yield showed the strongest positive trend among grade categories (r = 0.666, R² = 0.443, P = 0.334), consistent with the elevated Jumbo fractions observed under HT. Cull yield showed the strongest overall association with soil temperature (r = 0.851, R² = 0.724, P = 0.149), with HT recording the highest Cull yield (2.6 Mg ha^−^¹) across all treatments in 2023.

**Figure 5.**
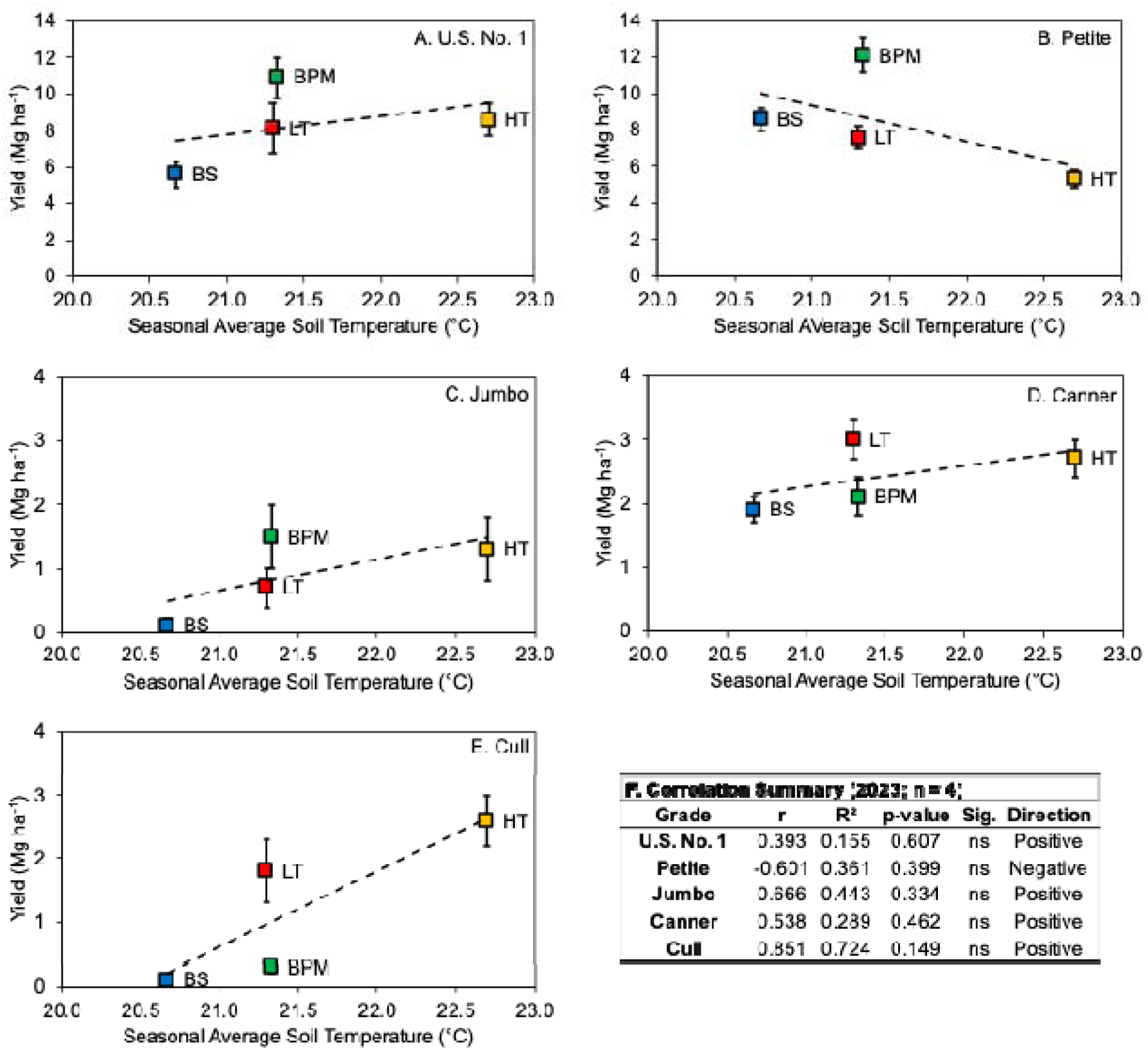
Relationship between seasonal average soil temperature (°C) at 12 cm depth and marketable yield (Mg ha^⁻^¹) by U.S. grade category for four protected culture treatments during the 2023 growing season at Rock Springs, PA. Panels A–E show individual grade categories: (A) U.S. No. 1, (B) Petite, (C) Jumbo, (D) Canner, and (E) Cull. Each data point represents one treatment: black plastic mulch (BPM), bare soil (BS), low tunnel (LT), and high tunnel (HT). Dashed lines represent linear regression fits for each grade. Panel F summarizes Pearson correlation coefficients (r), coefficients of determination (R²), p-values, and direction of association between seasonal average soil temperature and grade yield for all five grade categories (n = 4). Soil temperature was monitored using HOBO MX2202 pendant loggers (BPM, BS, and LT) and a Decagon EM50 datalogger with 5TM sensors (HT). Statistically significant correlations were detected at α = 0.05.

## DISCUSSION

### Yield responses to production system and season

Treatment effects on TY and MY were inconsistent across seasons, reflecting interactions between production system design and year-to-year weather variation. In 2022, HT recorded the highest TY (21.1 Mg ha^−^¹), while BS produced the highest MY (16.8 Mg ha^−^¹). In 2023, BPM was the top-performing treatment for both TY (26.8 Mg ha^−^¹) and MY (26.6 Mg ha^−^¹). These seasonal reversals underscore that no single system consistently dominated across both years, and that the yield advantage of any protected culture system depends on the prevailing thermal and moisture environment.

The yield range observed in this study is broadly consistent with prior sweetpotato production research in temperate and short-season regions. Duque (2020) reported MY values of 30.7 to 39.6 Mg ha^−^¹ for the highest-yielding cultivars grown under BPM at the same Rock Springs site, suggesting that the 2023 BPM yields observed here (26.6 Mg ha^−^¹) are within the range achievable under favorable conditions. Wees et al. (2015, 2016) reported a mean MY of approximately 24.4 Mg ha^−^¹ under BPM with short-season cultivars, similar to the BPM and BS values observed here in 2023. Sideman (2015) reported MY of 28.2 to 49.7 Mg ha^−^¹ for the best-performing cultivar-date combinations in New Hampshire, and Duque et al. (2022) reported a range of 4.4 to 70.7 Mg ha^−^¹ across planting configurations at the same site, illustrating the wide yield potential under optimized management.

The 2022 and 2023 seasons differed markedly in weather characteristics, with 2022 warmer and drier and 2023 cooler and wetter. These differences shaped the grade distribution: 2023 was characterized by substantially higher U.S. No. 1 and Petite yields and lower Jumbo and Cull fractions across all treatments compared with 2022. The elevated Jumbo and Cull yields in 2022, especially under HT, are consistent with excessively warm growing conditions and prolonged root development, which favor oversized or misshapen storage roots. Cull yield under HT was notably high in both seasons (4.8 Mg ha^−^¹ in 2022 and 2.6 Mg ha^−^¹ in 2023), suggesting that the high-tunnel environment may promote conditions that increase the proportion of unmarketable roots, particularly in warmer years.

### Effects of black plastic mulch and soil temperature

BPM is widely documented to raise soil temperature and improve sweetpotato yield in temperate regions by accelerating early-season root establishment (Duque, 2020; Wees et al., 2015, 2016; Sideman, 2015). In this study, BPM recorded a seasonal average soil temperature of 21.33°C in 2023, compared with 20.67°C under BS, a difference of 0.66°C. In a 3-year Manitoba study, Rao et al. (2022) reported that biodegradable black plastic mulch raised mean soil temperature by approximately 0.92°C (19.95 vs. 19.03°C at a depth of 5.1 cm), translating to a 3-year average MY increase of 39% over no-mulch. The modest soil temperature difference between BPM and BS at Rock Springs in 2023, combined with the large yield advantage of BPM (26.6 vs. 16.2 Mg ha^−^¹ for MY), suggests that the BPM benefit extends beyond soil warming alone and likely encompasses moisture retention, weed suppression, and protection against soil crusting during rainfall events (Amare and Desta, 2021).

Shrestha and Miles (2022) reported that polyethylene mulch raised soil temperature by approximately 3°C over bare ground in northwestern Washington, and biodegradable mulch by 2°C. LaHue et al. (2026) similarly reported that polyethylene mulch raised mean soil temperature by 1.8°C over bare ground at 10-cm depth in northwestern Washington and produced five to six times greater total and marketable yield than bare soil, underscoring that the thermal and agronomic benefits of plastic mulch are consistent across geographically distinct northern sweetpotato production environments. In the present study, no wireworm pressure was documented, and the yield advantage of BPM over BS was consistent in direction across both seasons, supporting the interpretation that BPM provided a reliable agronomic benefit under Pennsylvania conditions. Dinu et al. (2022) reported that polyethylene mulch increased total sweetpotato yield from a three-cultivar mean of 32.97 to 40.05 Mg ha^−^¹ across two seasons in Romania, a 21% increase, and improved root quality attributes, including total soluble solids, starch content, and vitamin C concentration.

Weed management represents an additional functional benefit of BPM that is difficult to quantify in yield terms alone. Laurie et al. (2015) demonstrated in South Africa that plastic mulch achieved greater than 90% canopy cover by week 5, effectively suppressing weeds throughout the season. In the present study, the BS treatment required supplemental cultivation and hand weeding before canopy closure, whereas BPM plots required minimal in-season weed management. This labor reduction is both agronomically and economically relevant for small-to medium-scale sweetpotato operations in Pennsylvania.

### High tunnel and low tunnel performance

The HT treatment offered the earliest transplant dates (13 May 2022 and 27 Apr 2023), the longest growing season, and the warmest and most stable soil temperatures (a seasonal average of 22.70 °C in 2023). Despite these thermal advantages, HT did not consistently produce the highest MY across both seasons, ranking first in TY in 2022 (21.1 Mg ha^−^¹) but third in both TY (20.4 Mg ha^−^¹) and MY (17.8 Mg ha^−^¹) in 2023. The elevated Cull fraction under HT in both seasons suggests that the high-tunnel environment may promote conditions that reduce root quality, potentially through elevated temperatures, restricted air circulation, or root congestion from extended vine growth.

Hoppenstedt et al. (2019) examined HT use for organic sweetpotato slip production in Kansas and found that HT provided higher slip yields in one of two seasons, suggesting context-dependent and season-sensitive benefits. That study reported mean HT soil temperatures of approximately 24-26°C during the bedding period, comparable to the peak temperatures recorded in the HT treatment in the present study. The authors noted that HT slip production may be a viable regional strategy for northern growers seeking to reduce their dependence on southern-produced slips, a consideration also relevant to Pennsylvania producers.

The LT treatment exhibited the most variable yield performance of the four systems. In 2022, LT produced the lowest TY (6.4 Mg ha^−^¹) and MY (5.1 Mg ha^−^¹) of any treatment, despite accumulating the second-highest seasonal GDD (618.8). This poor performance is attributed to establishment challenges, excessive humidity under the tunnel cover prior to removal, and possible vine disturbance during tunnel handling. In 2023, LT recovered substantially (TY: 21.2 Mg ha^−^¹; MY: 19.4 Mg ha^−^¹), more consistent with expectations for a season-extension system. The erratic LT response across seasons highlights the management sensitivity of low-tunnel systems: their agronomic benefit depends on timely installation and removal, adequate ventilation, and careful handling to avoid vine damage.

### Growing degree day accumulation and yield

Cumulative GDD (base 15.5°C) from transplanting to harvest ranged from 493.0 to 650.5 across treatments and years, well below the approximately 2,600 GDD typical of Louisiana production environments (Villordon et al., 2009). At the same Rock Springs site, Duque et al. (2022) identified a threshold of approximately 700 GDD (base 15.5°C) as the minimum for adequate Petite grade yield, a value approached but not always reliably exceeded in the present study, particularly in the cooler 2023 season (493 to 517 GDD across BPM, BS, and LT). HT accumulated the most GDD in both seasons (650.5 in 2022 and 516.6 in 2023) owing to its earlier transplant date, illustrating the practical value of season extension even in the absence of a large per-day temperature advantage.

The linear relationship between cumulative GDD and yield was negative and non-significant across both seasons (combined R² = 0.20 for TY, P = 0.266; R² = 0.38 for MY, P = 0.104), driven largely by anomalously low LT yields in 2022 despite moderate GDD accumulation. Villordon et al. (2009) demonstrated strong GDD-based harvest prediction models in Louisiana, where the relationship between thermal accumulation and root development is more consistent owing to longer and more uniform growing seasons. LaHue et al. (2026) demonstrated that soil GDD accumulation under polyethylene mulch and bare soil could each be estimated from air GDD using linear regression (R² = 0.99 in both cases) and concluded that a base temperature of 10°C captured continued root development and yield gains later in the season more accurately than the commonly used 15.5°C base temperature. In short-season temperate environments, GDD-based harvest scheduling serves primarily as a planning tool to estimate relative harvest readiness across treatments, rather than as a predictive model of absolute yield.

### Soil temperature and yield relationships

In 2023, the seasonal average soil temperature ranged from 20.67°C (BS) to 22.70°C (HT), with BPM (21.33°C) and LT (21.30°C) recording intermediate values. The relationship between seasonal average soil temperature and TY was weak and non-significant (R² = 0.038, P = 0.806), as was the relationship with MY (R² = 0.002, P = 0.958). These non-significant relationships reflect the small between-treatment range in soil temperature (approximately 2°C) and the confounding influence of other yield-determining factors. Soil temperature data were not available for 2022, precluding direct between-year comparison of soil thermal effects.

Rao et al. (2022) reported that even a modest mean soil temperature increase of approximately 1°C under BPM was associated with consistent MY gains of 30 to 47% across three seasons in Manitoba, demonstrating that small thermal differences can have agronomically significant yield consequences when integrated across the full growing season. In the present study, the 0.66°C soil temperature advantage of BPM over BS was accompanied by a yield advantage of 10.4 Mg ha^−^¹ MY in 2023, suggesting that non-thermal mechanisms of BPM, including moisture conservation, weed suppression, and physical protection of the root zone, contributed substantially to the observed yield advantage. LaHue et al. (2026) corroborated this finding by reporting a 1.8°C increase in soil temperature under polyethylene mulch in northwestern Washington, a cool maritime environment similar in thermal character to the humid-temperate conditions at Rock Springs, PA, thereby reinforcing the consistency of mulch-mediated soil warming across northern production regions.

### Cultivar performance

Cultivar response to treatment and season was highly variable, consistent with the broad range of genotypic adaptation reported among sweetpotato cultivars in temperate environments (Duque, 2020; Duque et al., 2022; Wees et al., 2015). In 2023, ’O’Henry’ under BPM recorded the highest TY (30.9 Mg ha^−^¹) and MY (30.8 Mg ha^−^¹) of any treatment-cultivar combination across both years. ’Beauregard’ and ’Averre’, consistently high performers in earlier Rock Springs cultivar trials (Duque, 2020; Duque et al., 2022), remained competitive under BPM, with 2023 TY values of 30.6 and 30.0 Mg ha^−^¹, respectively. ’Murasaki’ underperformed in 2023, recording the lowest TY under both BPM (19.0 Mg ha^−^¹) and LT (6.0 Mg ha^−^¹), consistent with its reputation as a cultivar better adapted to warmer, drier conditions. These results highlight the importance of cultivar selection as a key management decision, particularly in the context of production system choice, consistent with cultivar-by-system interactions reported in other northern sweetpotato production trials (LaHue et al., 2026).

## CONCLUSIONS

This study demonstrated that sweetpotato can be produced successfully in a temperate, short-season environment in central Pennsylvania across multiple production systems, with commercially relevant marketable yield (MY) achieved under black plastic mulch (BPM), bare soil (BS), high tunnel (HT), and low tunnel (LT) in at least one of the two seasons evaluated. No single production system dominated consistently across both years, and the yield advantage of any given system was contingent on seasonal weather conditions, particularly the prevailing temperature and moisture regime during early establishment.

BPM emerged as the most reliable system overall, producing the highest MY in 2023 (26.6 Mg ha^−^¹) and intermediate MY in 2022 (12.8 Mg ha^−^¹), while also providing weed suppression and moisture conservation benefits that reduced management inputs relative to BS. These findings are consistent with reports from other northern production environments, including northwestern Washington, where LaHue et al. (2026) reported five to six times greater total and marketable yield under polyethylene mulch than bare soil and a mean soil temperature increase of 1.8°C at 10-cm depth under mulch. For Pennsylvania growers considering sweetpotato adoption or system expansion, BPM represents the most agronomically consistent and practically accessible option, supported by a growing body of regional and national literature across diverse temperate production environments.

HT offered the earliest planting opportunity and the warmest soil environment, but these advantages did not consistently translate into the highest MY. The elevated Cull fractions observed in both seasons, particularly in 2022, suggest that high-tunnel conditions may limit root quality in warmer years. HT may be better suited to early-season transplant production, as a market differentiation strategy for early harvest, or for operations in which extended season length is the primary objective.

LT performance was highly variable, ranging from the lowest MY of any treatment in 2022 (5.1 Mg ha^−^¹) to a competitive value in 2023 (19.4 Mg ha^−^¹). The 2022 LT underperformance is attributed to management challenges related to tunnel ventilation and vine disturbance rather than a fundamental limitation of the system concept. Growers with experience managing low tunnels for other vegetable crops may find LT an accessible season-extension tool for sweetpotato, if installation, ventilation, and removal are managed carefully.

Cumulative GDD at this location remained substantially below the thresholds typical of major sweetpotato production regions, confirming that thermal limitation is the primary constraint for sweetpotato production in Pennsylvania. LaHue et al. (2026) demonstrated that a base temperature of 10°C captures continued root development and yield gains later in the season more accurately than the 15.5°C base commonly used in sweetpotato GDD models, a finding that warrants evaluation in future Pennsylvania trials. Matching cultivar selection to the production system, with attention to grade distribution targets and quality outcomes, represents an important additional lever for optimizing economic returns within these thermal constraints.

Future research should address the lack of 2022 soil temperature data by deploying monitoring equipment in both years of any subsequent trial. Additional evaluation of biodegradable mulch alternatives that reduce end-of-season plastic disposal concerns would support the sustainability objectives of Pennsylvania vegetable growers, building on results from Manitoba (Rao et al., 2022) and northwestern Washington (LaHue et al., 2026), where biodegradable mulch performed comparably to conventional polyethylene mulch in raising soil temperature and improving yield. Continued multi-season, multi-cultivar trials across protected culture systems will help refine production recommendations for the mid-Atlantic region, where sweetpotato acreage has the potential to expand as growers diversify operations and respond to growing regional demand for locally produced root vegetables.

## Acknowledgments

This work was supported by the Pennsylvania Department of Agriculture Specialty Crop Block Grant Program under Award Agreement C9400000527 and the U.S. Department of Agriculture, National Institute of Food and Agriculture, Hatch Appropriations under Project #PEN04582 and Accession #1005492. The author thanks the farm staff and student interns at the Russell E. Larson Agricultural Research Center, The Pennsylvania State University, for technical assistance and support. The author also thanks K.V. Pecota for providing thoughtful advice throughout this project and acknowledges the National Sweetpotato Collaborators Group for collegial advice and collaboration.

